# Rice paddy *Nitrospirae* encode and express genes related to sulfate respiration: proposal of the new genus *Candidatus* Sulfobium

**DOI:** 10.1101/196774

**Authors:** Sarah Zecchin, Ralf C. Mueller, Jana Seifert, Ulrich Stingl, Karthik Anantharaman, Martin van Bergen, Lucia Cavalca, Michael Pester

**Author notes:** contributed equally (Sarah Zecchin, Ralf C. Mueller). Corresponding author: Michael Pester, Department Microorganisms, Leibniz Institute DSMZ — German Collection of Microorganisms and Cell Cultures, D-38124 Braunschweig, Germany.

## Abstract

*Nitrospirae* spp. distantly related to thermophilic, sulfate-reducing *Thermodesulfovibrio* species are regularly observed in environmental surveys of anoxic marine and freshwater habitats. However, little is known about their genetic make-up and physiology. Here, we present the draft genome of *Nitrospirae* bacterium Nbg-4 as a representative of this clade and analyzed its *in situ* protein expression under sulfate-enriched and sulfate-depleted conditions in rice paddy soil. The genome of Nbg-4 was assembled from replicated metagenomes of rice paddy soil that was used to grow rice plants in the presence and absence of gypsum (CaSO_4_×2H_2_O). Nbg-4 encoded the full pathway of dissimilatory sulfate reduction and showed expression thereof in gypsum-amended anoxic bulk soil as revealed by parallel metaproteomics. In addition, Nbg-4 encoded the full pathway of dissimilatory nitrate reduction to ammonia, which was expressed in bulk soil without gypsum amendment. The relative abundance of Nbg-4-related metagenome reads was similar under both treatments indicating that it maintained stable populations while shifting its energy metabolism. Further genome reconstruction revealed the potential to utilize butyrate, formate, H_2_, or acetate as electron donor, with the Wood-Ljungdahl pathway being expressed under both conditions. Comparison to publicly available *Nitrospirae* genome bins confirmed that the pathway for dissimilatory sulfate reduction is also present in related *Nitrospirae* recovered from groundwater. Subsequent phylogenomics showed that such microorganisms form a novel genus within the phylum *Nitrospirae*, with Nbg-4 as a representative species. Based on the widespread occurrence of this novel genus, we propose for Nbg-4 the name *Candidatus* Sulfobium mesophilum, gen. nov., spec. nov.

**Importance:** Rice paddies are indispensable for food supply but are a major source of the greenhouse gas methane. If not counterbalanced by cryptic sulfur cycling, methane emission from rice paddy fields would be even higher. However, the microorganisms involved in this sulfur cycling are little understood. By using an environmental systems biology approach of Italian rice paddy soil, we could retrieve the population genome of a novel member of the phylum *Nitrospirae*. This microorganism encoded the full pathway of dissimilatory sulfate reduction and expressed it *in situ* under sulfate-enriched and anoxic conditions. Phylogenomics and comparison to environmental surveys showed that such microorganisms are actually widespread in freshwater and marine environments. At the same time, they represent a yet undiscovered genus within the little explored *Nitrospirae*. Our results will be important to design enrichment strategies and postgenomic studies to fully understand the contribution of these novel *Nitrospirae* to the global sulfur cycle.

## Introduction

Sulfate reducing microorganisms (SRM) are regularly observed in rice paddy fields (1-8). Despite the prevailing low sulfate concentrations in this habitat (lower μM-range, 9, 10), the rice rhizosphere and bulk soil are characterized by high sulfate reduction rates, which are comparable to marine surface sediments (11). This at first sight contradictory observation is explained by a cryptic sulfur cycle. Here, the small sulfate pool is rapidly reduced to sulfide but the latter also rapidly re-oxidized to sulfate thus keeping a highly active sulfur cycling running (10-13). This cryptic sulfur cycle can occur at oxic-anoxic interfaces such as rice roots but apparently runs also in the completely anoxic bulk soil (10). Under the latter conditions, reduced sulfur species may be re-oxidized with the help of iron minerals or redox-active parts of humic material such as quinone moieties as shown for other freshwater habitats (14-16).

The ability to perform dissimilatory sulfate reduction is most widespread among members of the *Deltaproteobacteria* and *Firmicutes* (17). Additional and exclusively thermophilic sulfate reducers are affiliated to the archaeal phyla *Euryarchaeota* and *Crenarchaeota* and the bacterial phyla *Thermodesulfobacteria* and *Nitrospirae* (17, 18). The only known SRM in the phylum *Nitrospirae* are bacteria belonging to the genus *Thermodesulfovibrio* (19-23). All described species of this genus are thermophilic with their common metabolic properties comprising the reduction of sulfate, thiosulfate and in some cases sulfite with a limited range of electron donors. These include pyruvate and lactate, which are incompletely oxidized to acetate, or H_2_ and formate in a background of acetate as auxiliary carbon source. Especially the inability for autotrophic growth and the incomplete oxidation of organic substrates to acetate is a characteristic feature of this genus. Alternative electron acceptors used by *Thermodesulfovibrio* spp. are Fe(III) and in the case of *Thermodesulfovibrio islandicus* DSM 12570 nitrate (19-23).

In addition to the genus *Thermodesulfovibrio*, the phylum *Nitrospirae* currently encompasses the genera *Nitrospira* and *Leptospirillum*, which comprise species exclusively involved in nitrification or iron reduction, respectively (24, 25). A group of still uncultured *Nitrospirae*, which form a sister clade to the genus *Thermodesulfovibrio*, is represented by magnetotactic bacteria belonging to the putative genera *Candidatus* Magnetobacterium (26-28), *Candidatus* Thermomagnetovibrio (29), *Candidatus* Magnetoovum (30, 31), and *Candidatus* Magnetominusculus (32). These microorganisms are typically encountered at the oxic-anoxic interface of sediments but were also enriched from water of hot springs (33). The observation of sulfur-rich inclusions in the cells of *Ca*. Magnetobacterium bavaricum (27), *Ca*. Magnetoovum chiemensis (31), and Ca. Magnetoovum mohavensis (30), the presence of sulfur metabolism genes in the genomes of the former two species (31), and their predominant occurrence at oxic-anoxic interfaces led to the hypothesis that these microorganisms could be involved in sulfur oxidation (27, 31, 33).

All SRM have the canonical pathway of dissimilatory sulfate reduction in common, which is an intracellular process that involves an eight-electron reduction of sulfate to sulfide. This pathway proceeds through the enzymes sulfate adenylyltransferase (Sat), adenylyl posphosulfate reductase (Apr), dissimilatory sulfite reductase (Dsr), and the sulfide-releasing DsrC (34). In addition, the complexes QmoAB(C) and DsrMK(JOP) are important in transferring reducing equivalents towards the pathway of sulfate reduction (35). The only known exception to this rule are ANME-archaea that anaerobically oxidize methane by a yet unresolved mechanism of sulfate reduction to zero-valent sulfur (36). The two different subunits of the heterotetrameric dissimilatory sulfite reductase Dsr are encoded by the paralogous genes *dsrA* and *dsrB*, which are frequently used as functional phylogenetic markers for SRM (37). The phylogeny of reductive bacterial-type DsrAB is subdivided into the *Deltaproteobacteria*, *Firmicutes*, Environmental, and *Nitrospirae* superclusters (37). DsrAB sequences affiliated with the *Nitrospirae* supercluster were predominantly found in freshwater and soil environments and to a smaller extent in marine, industrial, or hot-temperature habitats (37). Intriguingly, they were also detected before in Italian (10) and Chinese (4, 8) rice paddy soils, but the detailed phylogenetic affiliation of these *dsrAB*-carrying microorganisms and their possible involvement in rice paddy sulfur cycling remained unclear.

Here, the draft genome of a novel and putatively sulfate reducing species belonging to the phylum *Nitrospirae* has been obtained from a metagenome survey of rice paddy soil. We present its metabolic potential and phylogeny as reconstructed from its genome and compare this to *Nitrospirae* genome bins recently recovered from metagenome studies of groundwater habitats. To support our conclusions, we present *in situ* protein expression patterns of this novel *Nitrospirae* species as inferred by a metaproteome analysis of rice paddy soil.

## Results

### A *Nitrospirae* genome from rice paddy soil

We used a metagenomics approach to identify novel microorganisms involved in rice paddy sulfur cycling. For this purpose, replicated metagenomes (Table S1) were sequenced from bulk and rhizosphere soils of rice plants, which were grown either in gypsum-amended (CaSO_4_×2H_2_O) or un-amended (control) soils. Among the 159 population genome bins that could be retrieved, *Nitrospirae* genome bin Nbg-4 was outstanding because it encoded *dsrAB*, was of high quality with ≤2% residual contamination, showed no strain heterogeneity, and had an estimated genome completeness of 75% (Table 1). The relative abundance of Nbg-4 was highest in the bulk soils averaging 17 RPKM (reads per kilobase of scaffold per million reads) and roughly three times lower in rhizosphere soils (Figure 1). A two-way analysis of variance (ANOVA) showed that soil compartment had a significant effect on the relative abundance of Nbg-4 (F_2,14_=36.16, p<0.001), while gypsum amendment (F_1,14_=0.17, p=0.69) and the interaction of soil compartment and gypsum amendment (F_1,12_=0.03, p=0.87) remained insignificant. To estimate the index of replication (iRep, 38) of Nbg-4, single reads of metagenomic replicates were combined per soil habitat to achieve sufficient coverage. This analysis indicated that roughly three quarters of the population were replicating their genome in freshly flooded soils, while roughly one third replicated its genome in bulk soils after 58-59 days of incubation irrespective of gypsum treatment (Table 1). For rhizosphere soils, the coverage was not sufficient to perform an iRep analysis.

**Table 1.** Characteristics of the obtained draft genome of *Nitrospirae* bacterium Nbg-4.

| *Nitrospirae* bacterium Nbg-4 |  |
| --- | --- |
| **Genome feature** |  |
| Chromosome size (Mbp) | 2.77 |
| GC content (%) | 49 |
| Number of scaffolds | 151 |
| Number of CDS | 2855 |
| Average CDS length (bp) | 855 |
| Protein coding density (%) | 87 |
| Number of rRNA genes | 1 |
| Number of tRNA genes | 21 |
| **ChekM analysis** |  |
| Completeness (%) | 75.5 |
| Contamination (%) | 2.0 |
| Strain heterogeneity (%) | 0.0 |
| **iRep analysis** |  |
| iRep initial soil | 1.73 |
| iRep bulk soil without gypsum | 1.34 |
| iRep bulk soil with gypsum | 1.31 |

**Figure 1.**
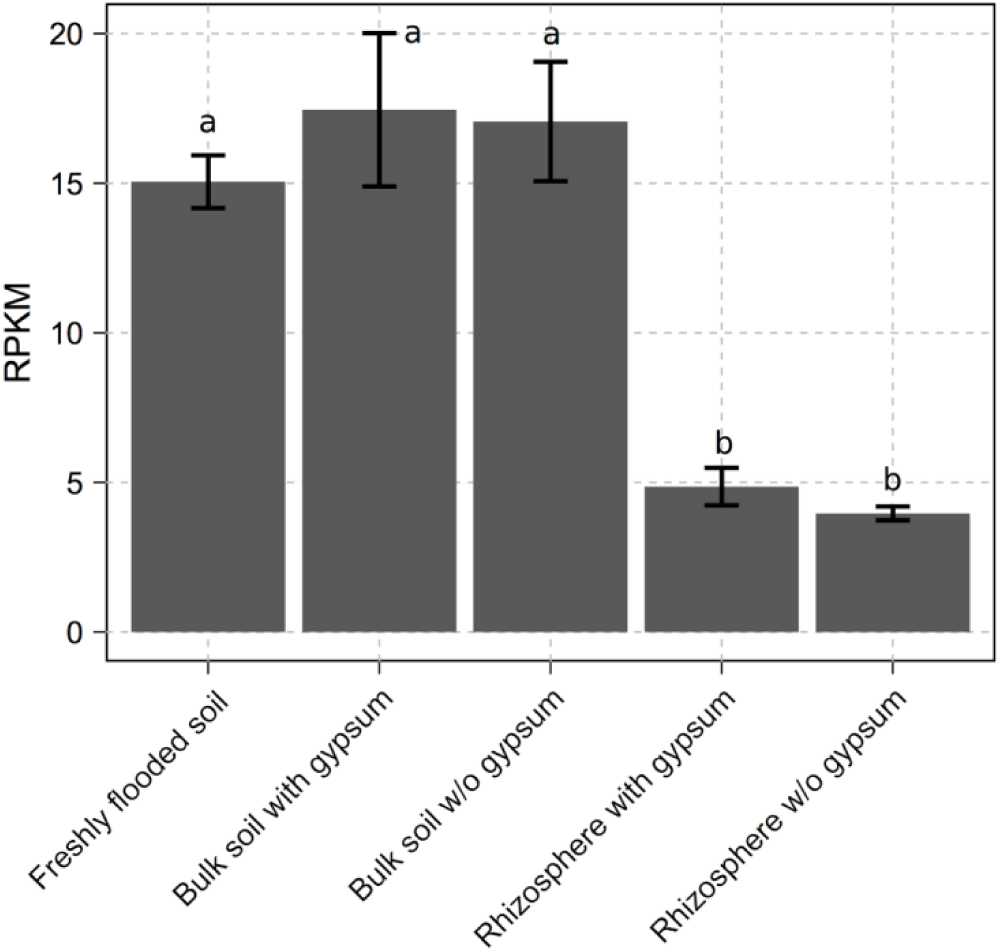
Average relative abundance (± one standard deviation) of *Nitrospirae* bacterium Nbg-4 in the differently treated soil habitats as inferred from the RPKM values (reads per kilobase of scaffold per million reads) of its longest scaffold. Significant differences are indicated by different letters and were inferred by a two-way ANOVA and a post-hoc Tukey test (p<0.001).

### Reconstruction of a dissimilatory sulfur metabolism

The complete pathway for dissimilatory sulfate reduction was recovered in Nbg-4 (Figure 2). Besides genes encoding Sat and the β-subunit of Apr, which catalyze the activation of sulfate and its concomitant reduction to sulfite, respectively, also genes for DsrAB and DsrC, which reduce sulfite further to sulfide could be detected. *aprA* was missing because of an assembly break in the scaffold after *aprB* (typically *aprA* is downstream of *aprB*). In addition, genes encoding the electron-transferring QmoABC and DsrMK were detected. *Thermodesulfovibrio* spp. possess in addition to the module DsrMK also the module DsrJOP, which form together the membrane-bound electron-transferring complex DsrMKJOP (23, 35). Since *dsrMK* were located at the end of one scaffold in Nbg-4 and another scaffold started with a long fragment of *dsrP*, it is likely that also Nbg-4 encodes a complete DsrMKJOP complex. In support of a reductively operating sulfur metabolism, the presence of *dsrD* directly adjacent to *dsrAB* was detected. DsrD is a small protein of putative regulatory function present in all sulfate reducers (39) with sporadic encounters in genomes of sulfide and sulfur-oxidizing bacteria (40). In addition, *dsrN* and *dsrT* as typical genes of the dsr operon in sulfate reducers and sulfur-oxidizing green sulfur bacteria (39, 41) and *hppA*, which codes for a membrane-bound and proton-translocating pyrophosphatase to pull, e.g., the energy-demanding reaction of Sat, were detected.

**Figure 2.**
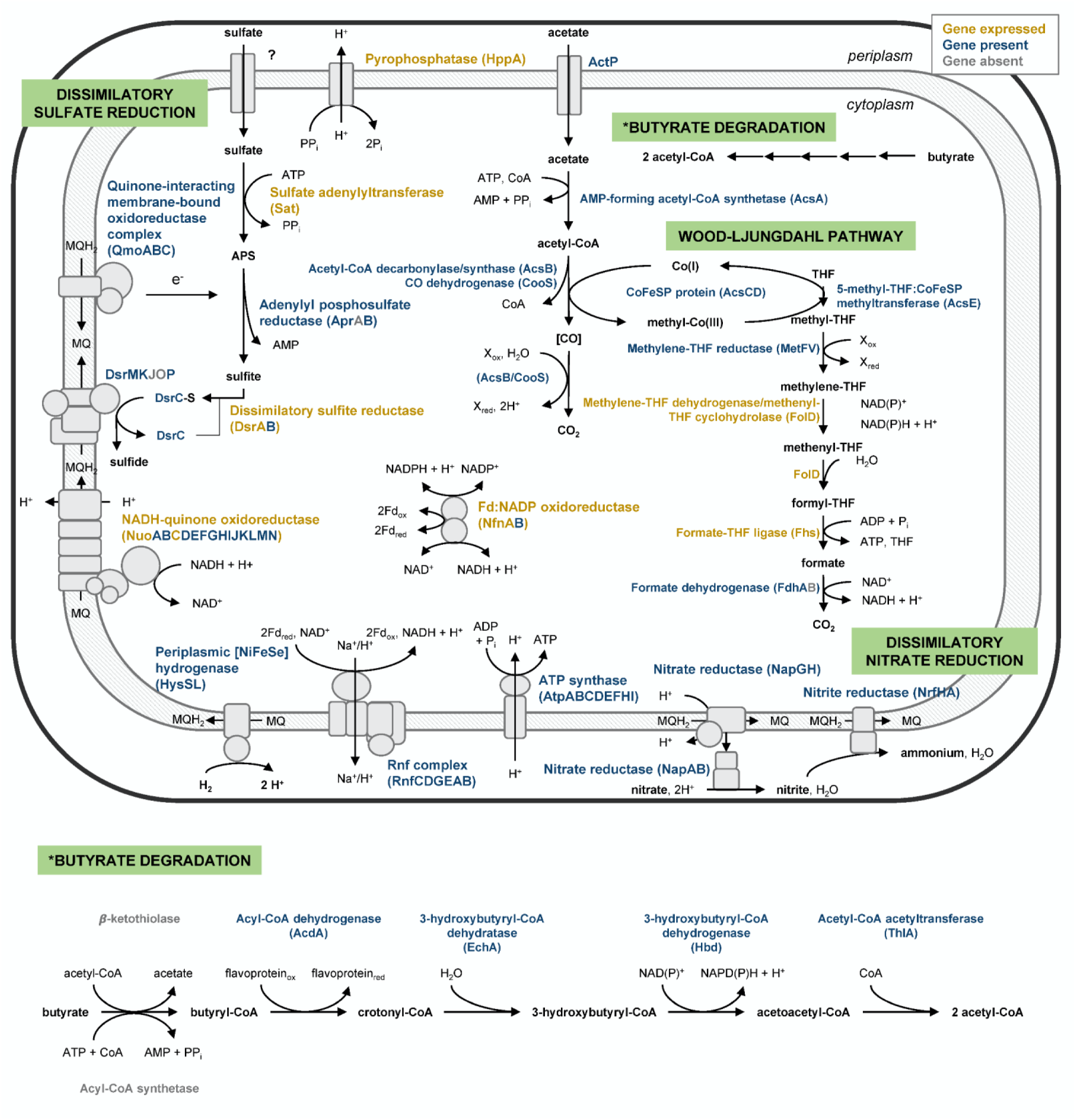
Schematic view of reconstructed energy metabolism pathways in *Nitrospirae* bacterium Nbg-4. *In situ* expression of proteins in bulk soil treated with gypsum as revealed by metaproteomics is color-indicated. Protein expression in other soil habitats and treatments is given in Table S2.

All soil samples that were used for metagenome sequencing were also analyzed for their metaproteome. In bulk soil treated with gypsum, a search against Nbg-4 encoded proteins identified peptides specific for Sat and DsrA as two essential components of the first and last step of sulfate reduction, respectively (Figure 2). Peptides specific for DsrA of Nbg-4 were also detected in rhizosphere soil treated with gypsum. In contrast, no peptides matching Nbg-4 sulfur metabolism proteins were detected in control treatments without gypsum, neither in the bulk soil nor in the rhizosphere (Table S2). The fragmented recovery of proteins involved in dissimilatory sulfate reduction is certainly a result of the low coverage of the proteome of a single microbial population in the background of the whole soil metaproteome.

Based on the recovery of the dissimilatory sulfate reduction pathway in Nbg-4, NCBI’s sequence repositories were searched for additional *dsrAB*-carrying *Nitrospirae* genome bins of high assembly quality. This analysis identified fourteen additional bins recovered from metagenomes: three from aquifer sediments (42), nine from aquifer groundwater (42), and two from a deep subsurface water (43) (Table S3). In-depth analysis of four bins that represent the three additional habitat types revealed not only the presence of *dsrAB* but also of the complete dsr operon including *dsrC*, *dsrD*, *dsrN*, *dsrT*, and *dsrMKJOP*, which were all in synteny to the respective genes of Nbg-4 (Figure. 3). Only *Nitrospirae* bacterium CG1-02-44-142 recovered from deep subsurface water had an inversion of *dsrC*, *dsrT*, and *dsrMKJOP* on its genome. Interestingly, also all other components of the dissimilatory sulfate reduction pathway including *sat*, *aprBA*, *qmoABC*, and *hppA* were encoded on these *Nitrospirae* genome bins, either completely or partially depending on the assembly breaks of the respective scaffolds (Table 2).

**Table 2.**
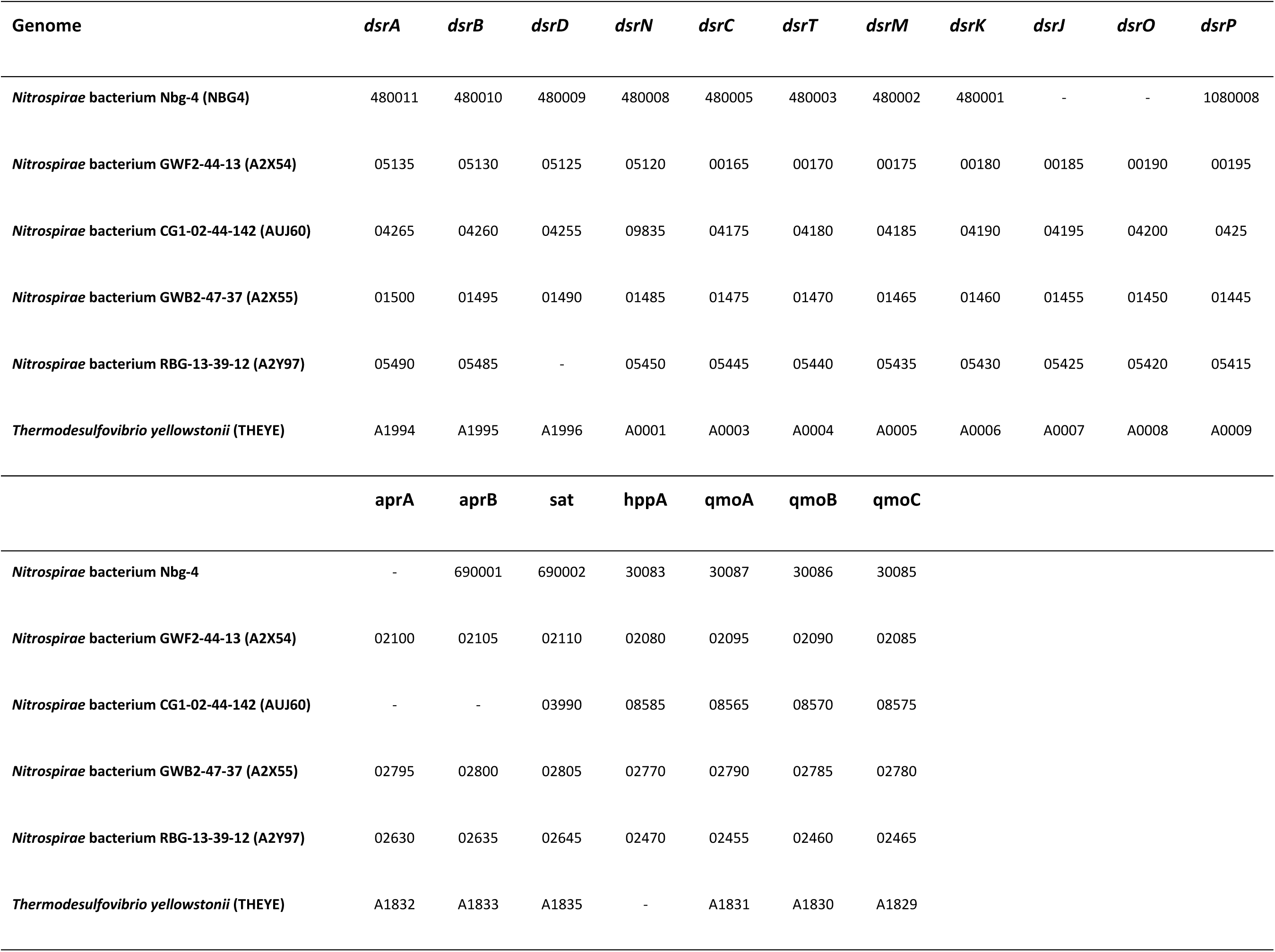
Locus tag of genes involved in dissimilatory sulfate reduction in *Nitrospirae* bacterium Nbg-4, related *dsrAB*-carrying *Nitrospirae* recovered from groundwater metagenomes (42, 43), and *Thermodesulfovibrio yellowstonii*.

| Genome | *dsrA* | *dsrB* | *dsrD* | *dsrN* | *dsrC* | *dsrT* | *dsrM* | *dsrK* | *dsrJ* | *dsrO* | *dsrP* |
| --- | --- | --- | --- | --- | --- | --- | --- | --- | --- | --- | --- |
| *Nitrospirae* bacterium Nbg-4 (Nbg4) | 480011 | 480010 | 480009 | 480008 | 480005 | 480003 | 480002 | 480001 | - | - | 1080008 |
| *Nitrospirae* bacterium GWF2-44-13 (A2X54) | 05135 | 05130 | 05125 | 05120 | 00165 | 00170 | 00175 | 00180 | 00185 | 00190 | 00195 |
| *Nitrospirae* bacterium CG1-02-44-142 (AUJ60) | 04265 | 04260 | 04255 | 09835 | 04175 | 04180 | 04185 | 04190 | 04195 | 04200 | 0425 |
| *Nitrospirae* bacterium GWF2-47-37 (A2X55) | 01500 | 01495 | 01490 | 01485 | 01475 | 01470 | 01465 | 01460 | 01455 | 01450 | 01445 |
| *Nitrospirae* bacterium RBG-13-39-12 (A2Y97) | 05490 | 05485 | - | 05450 | 05445 | 05440 | 05435 | 05430 | 05425 | 05420 | 05415 |
| *Thermodesulfovibrio yellowstonii* (THEYE) | A1994 | A1995 | A1996 | A0001 | A0003 | A0004 | A0005 | A0006 | A0007 | A0008 | A0009 |
|  | *aprA* | *aprB* | *sat* | *hppA* | *qmoA* | *qmoB* | *qmoC* |  |  |  |  |
| *Nitrospirae* bacterium Nbg-4 | - | 690001 | 690002 | 30083 | 30087 | 30086 | 30085 |  |  |  |  |
| *Nitrospirae* bacterium GWF2-44-13 (A2X54) | 02100 | 02105 | 02110 | 02080 | 02095 | 02090 | 02085 |  |  |  |  |
| *Nitrospirae* bacterium CG1-02-44-142 (AUJ60) | - | - | 03990 | 08585 | 08565 | 08570 | 08575 |  |  |  |  |
| *Nitrospirae* bacterium GWF2-47-37 (A2X55) | 02795 | 02800 | 02805 | 02770 | 02790 | 02785 | 02780 |  |  |  |  |
| *Nitrospirae* bacterium RBG-13-39-12 (A2Y97) | 02630 | 02635 | 02645 | 02470 | 02455 | 02460 | 02465 |  |  |  |  |
| *Thermodesulfovibrio yellowstonii* (THEYE) | A1832 | A1833 | A1835 | - | A1831 | A1830 | A1829 |  |  |  |  |

**Figure 3.**
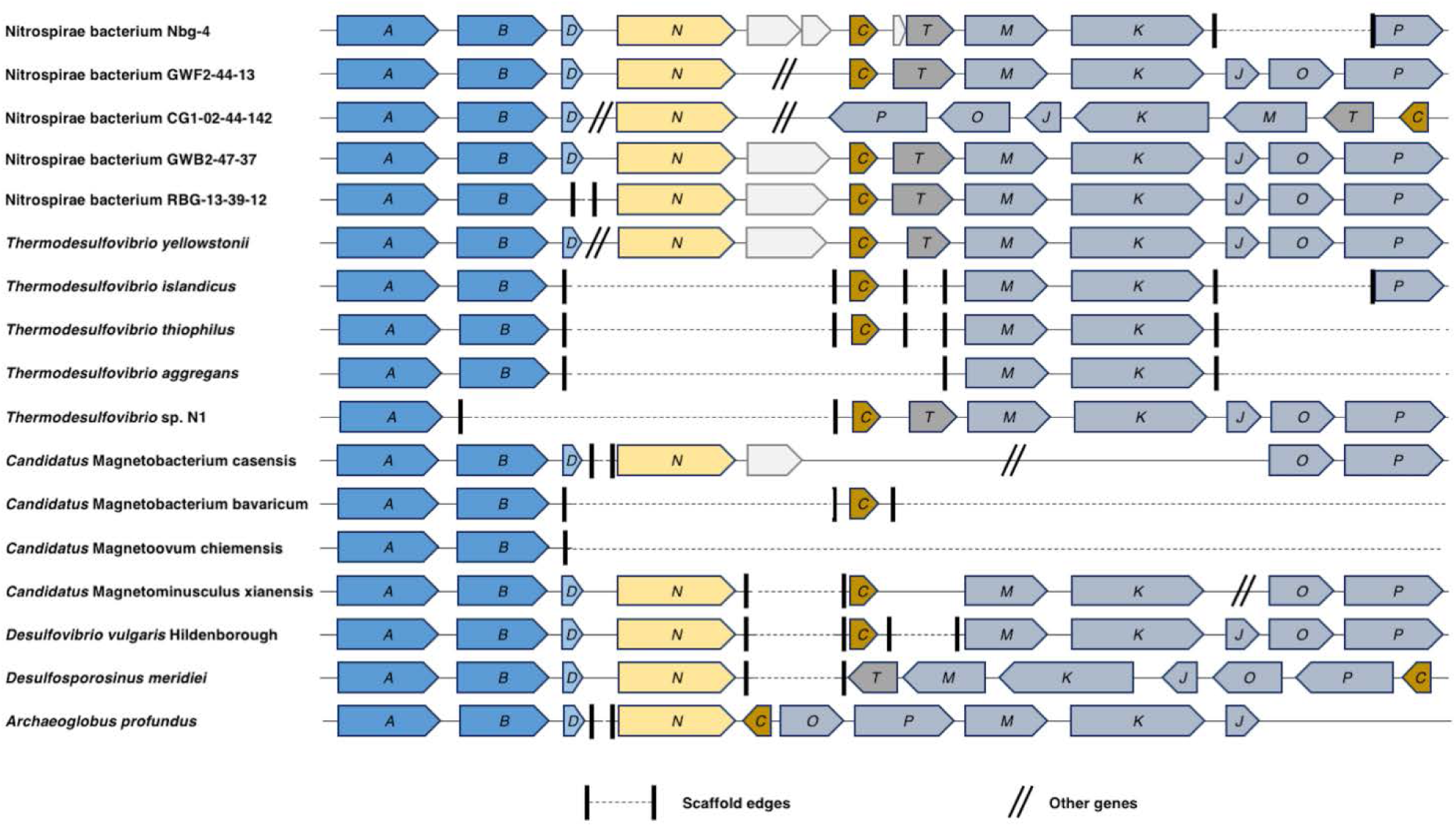
Organization and synteny of the dsr operon in *Nitrospirae* bacterium Nbg-4 in comparison to other *dsrAB*-carrying members of the phylum *Nitrospirae*. In addition, typical representatives of known sulfate-reducing microorganisms within the Deltaproteobacteria (*Desulfovibrio vulgaris* Hildenborough), Firmicutes (*Desulfosporosinus meridiei*), and Archaea (*Archaeoglobus fulgidus*) are shown.

### Nitrate reduction as an alternative respiratory metabolism

Nbg-4 also encoded a full set of genes necessary for dissimilatory nitrate reduction to ammonia (DNRA) (Figure 2). DNRA is employed by members of the genera *Thermodesulfovibrio*, *Desulfovibrio*, *Desulfobulbus*, *Desulfobacterium*, and *Desulfotomaculum* as alternative respiratory pathway in the absence of sulfate (39). The first step of DNRA is the reduction of nitrate to nitrite. To perform this step, Nbg-4 contains a periplasmic nitrate reductase NapA that forms a soluble complex with cytochrome c-containing NapB and couples electron transfer from the quinone pool by the membrane-associated quinol dehydrogenase module formed by NapGH (Table S2). In Nbg-4, the nap operon lacks NapC, which is a proposed electron-transferring, membrane-associated protein typically observed in DNRA-performing SRM. The lack of NapC resembles the situation in *Wolinella succinogenes* that also lacks this protein while being able to perform DNRA (44). The second step of DNRA employs a six-electron transfer to reduce nitrite to ammonia. In Nbg-4, this step is encoded by the membrane-bound nitrite reductase complex formed by NrfA, a periplasmic nitrite reductase, and NrfH, a membrane-associated quinol reductase that delivers electrons to NrfA. Screening of the obtained metaproteomes for DNRA-related proteins of Nbg-4, identified peptides specific for NapA and NapG in bulk soils without gypsum treatment. This indicates DNRA-activity of Ngb-4 under sulfate-depleted conditions. No expression of DNRA-related proteins was detected in bulk soil treated with gypsum or in the rhizosphere samples, irrespective of gypsum treatment (Table S2).

### The genetic potential for complete oxidation of organic matter to CO_2_

The genome of Nbg-4 encoded the capacity for complete oxidation of acetate to CO_2_. This included the acetate transporter ActP, activation of acetate to acetyl-CoA by an AMP-forming acetyl-CoA synthetase (AcsA) and the complete Wood-Ljungdahl pathway (Figure 2, Table S2). Peptides specific for several of these enzymes could be detected by metaproteomics both in the bulk soil and rhizosphere irrespective of gypsum treatment (Table S2). The Wood-Ljungdahl pathway included at the end of its methyl branch a formate dehydrogenase, which provides Nbg-4 with the potential to utilize also formate as an electron donor. In addition, a periplasm-oriented, membrane-bound [NiFeSe] hydrogenase (HysLS) was detected, which connects to the quinone pool in the membrane (Figure 2). However, no peptides related to either one of these two enzyme complexes could be detected (Table S2). Furthermore, the potential for butyrate degradation via a β-oxidation was encoded. With the exception of the activation step of butyrate to butyryl-CoA, all genes encoding for the necessary enzymes were recovered (Figure 2). Peptides that match Nbg-4 enzymes involved in butyrate degradation were detected in rhizosphere but not in bulk soil metaproteomes (Table S2).

Coupling of electron transfer to energy conservation could be mediated in Nbg-4 by an electron-bifurcating Fd:NADP oxidoreductase (NfnAB), a H^+^/Na^+^-pumping Rnf complex (RnfCDGEAB), and a NADH-quinone oxidoreductase (respiratory complex I, NuoABCDEFGHIJKLMN) (35). In addition, the full set of genes encoding the ATP synthase was identified (AtpABCDEFHI) (Figure 2). Peptides specific for each of these Nbg-4 enzyme complexes were identified in the various bulk and rhizosphere soil metaproteomes (Table S2), indicating their active role in electron transfer and energy conservation.

### Phylogenetic affiliation of the *Nitrospirae* genome bin Nbg-4

A phylogenomic maximum-likelihood tree placed Nbg-4 and eight of the fourteen *dsrAB*-carrying *Nitrospirae* bacteria recovered in other studies (Table 2) in a stable cluster that branched off between *Thermodesulfovibrio* spp. and magnetotactic *Nitrospirae*. Two additional *dsrAB*-carrying *Nitrospirae* bacteria (GWA2-46-11 and GWB2-47-37) formed a sister branch to the Nbg-4 containing cluster and were more closely related to *Thermodesulfovibrio* species (Figure 4a). The remaining four *dsrAB-*carrying *Nitrospirae* bacteria branched off more basely within the phylum Nitrospirae forming two separate lineages with no clear affiliation to previously isolated species (Figure 4a).

**Figure 4.**
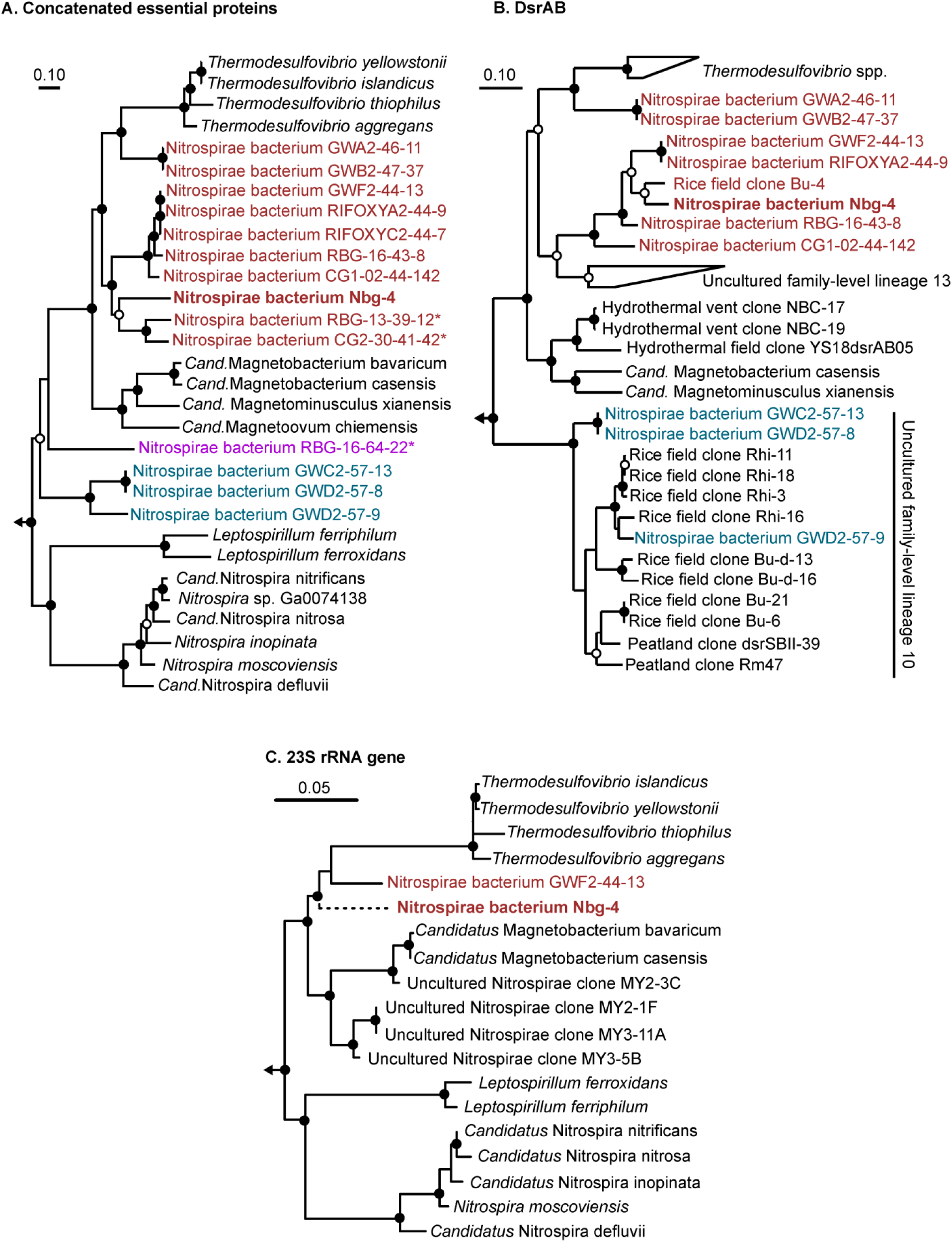
Phylogeny of *Nitrospirae* bacterium Nbg-4 (in bold) and related *dsrAB*-carrying *Nitrospirae* bacteria recovered from metagenomes of groundwater systems (42, 43). Uncultured *dsrAB*-carrying *Nitrospirae* bacteria that form separate genera as inferred by the genome-wide AAI approach are color coded. Maximum likelihood trees were inferred using the RAxML algorithm (71) and (A) a concatenated alignment of 43 essential proteins (Table S6), (B) deduced DsrAB sequences, and (C) the 23S rRNA gene. The partially recovered 23S rRNA gene of Nbg-4 was added to a RAxML tree of almost full-length 23S rRNA genes using the Quick add parsimony tool as implemented in ARB (74) without changing the tree topology. This is indicated by the dashed branch leading to Nbg-4 in this tree. Bootstrap support is indicated by closed (≥90%) and open (≥70%) circles at the respective branching points. The scale bar indicates 10 or 5% estimated sequence divergence, respectively.

The same branching pattern was recovered when analyzing deduced DsrAB sequences. Here, the well separated Nbg-4 containing cluster was most closely related to uncultured *dsrAB* family-level lineage 13 as defined by A. L. Müller et al. (37). Both clusters shared a common origin branching off between *Thermodesulfovibrio* species and magnetotactic *Nitrospirae* (Figure 4b). As in the phylogenomics appraoch, *Nitrospirae* bacteria GWA2-46-11 and GWB2-47-37 formed a stable sister branch that was more closely related to *Thermodesulfovibrio* species. Interestingly, the *dsrAB* of *Nitrospirae* bacterium RBG-13-39-12 and CG2-30-41-42, which were the closest relatives to Nbg-4 in the phylogenomics approach, did not fall into the *Nitrospirae* supercluster but were most closely related to uncultured *dsrAB* family-level lineage 11, which belongs to the Deltaproteobacteria supercluster (Fig. S1). This indicates lateral gene transfer of *dsrAB* within the phylum *Nitrospirae*, which is further supported by the DsrAB phylogeny of the basely branching Nitrospirae bacterium RBG-16-64-22. Here, the respective DsrAB sequences were clearly affiliated to the oxidative bacterial-type DsrAB having the alphaproteobacterium *Magnetococcus marinus* and *Chlorobi* spp. as closest relatives (Fig. S1). In contrast, DsrAB of *Nitrospirae* bacteria that formed the second basely branching lineage in the phylogenomics approach were also clustering basely in the DsrAB *Nitrospirae* supercluster and clustered within or as closest relatives to uncultured *dsrAB* family-level lineage 10 (Fig. 4B).

In a third approach, the phylogenetic position of the partial 23S rRNA gene of Nbg-4 was inferred when placed into a full-length 23S rRNA gene tree of cultured and uncultured members of the phylum *Nitrospirae*. Also here, Nbg-4 branched off between stable clusters related to *Thermodesulfovibrio* species and magnetotactic *Nitrospirae* (Figure 4c), thus corroborating the phylogenetic placement of the other two approaches.

In parallel, a genome-wide average nucleotide identity (gANI) and average amino acid identity (gAAI) analysis was performed (45-47). The gANI analysis revealed that all *Nitrospirae* genomes used for the phylogenomic tree reconstruction were less similar than 70% to the genome of Nbg-4 (Table S4). Since this is well below the proposed value of 96.5% to group bacterial strains into the same species (46), Nbg-4 represents a novel species. The gAAI analysis mainly mirrored the phylogenomic tree reconstruction. Here, all genomes within the Nbg-4 containing cluster as well as the sister branch that encompasses *dsrAB*-carrying *Nitrospirae* bacteria GWA2-46-11 and GWB2-47-37 shared identities between 55 and 100% (Table S5). At the same time, these genomes shared less than 55% identity to representatives of other genera within the *Nitrospirae*. In addition, the two basely branching lineages of *dsrAB*-carrying *Nitrospirae* genome bins represented either by *Nitrospirae* bacterium RBG-16-64-22 or *Nitrospirae* bacteria GWC2-57-13, GWD2-57-8, and GWD2-57-9 shared less than 55% gAAI identity to *Nitrospirae* spp. outside of their respective lineage. At the same time, the later three *dsrAB*-carrying *Nitrospirae* bacteria shared among themselves gAAI identities of 62-99% (Table S5). Since 55% gAAI is the lower boundary that is currently recommended to group bacterial strains into the same genus (45), Nbg-4 and the additional uncultured *dsrAB*-carrying *Nitrospirae* bacteria listed in Table S3 form three independent genera.

## Discussion

Members of the phylum *Nitrospirae*, which form a stable clade between thermophilic *Thermodesulfovibrio* spp. and magnetotactic *Nitrospirae*, are regularly observed in 16S rRNA gene- and *dsrAB*-based surveys of anoxic freshwater and marine environments of moderate temperature. These environments include marine (37) and estuarine (48) sediments, groundwater (42, 49), lake sediment (50), wetland soil (51), and rice paddy fields (10, 52, 53). Also in rice paddy soil analyzed in this study, eight species-level operational taxonomic units (OTUs) of such *Nitrospirae* were observed previously by 16S rRNA gene-based amplicon sequencing (Fig. S2, 7). So far, the genetic make-up and physiological characteristics of these microorganisms were largely unknown. Here, we present a detailed genome analysis of *Nitrospirae* bacterium Nbg-4 as a representative of this clade and analyzed its *in situ* protein expression profile under sulfate-enriched and sulfate-depleted conditions.

Nbg-4 encoded the complete pathway for dissimilatory sulfate reduction (Figure 2). Indeed, there are several lines of evidence that this newly discovered member of the *Nitrospirae* could represent an active sulfate reducer in rice paddy soil. From a genomic perspective, Nbg-4 carries not only all genes necessary for sulfate reduction but also genes of unknown function that are typically found in SRM such as *dsrD*, *dsrN* and *dsrT* (39). The same dsr operon organization (Figure 3) as well as the presence of all sulfate reduction-related genes (Table 2) were observed in the genomes of the other *dsrAB-*carrying *Nitrospirae* bacteria that form a stable phylogenetic lineage with Nbg-4 (Figure 4). From a phylogenetic perspective, *dsrAB* of Nbg-4 and related *Nitrospirae* bacteria were clearly affiliated to 11 the branch of reductively operating DsrAB of bacterial origin, which are phylogenetically separated from oxidatively operating DsrAB of bacterial origin (37). Most importantly, from an activity perspective the expression of enzymes involved in sulfate reduction was preferentially detected in gypsum-treated bulk soil, i.e. under completely anoxic and sulfate-enriched conditions. On the contrary, under sulfate-depleted conditions in control bulk soil, the expression of DNRA-related enzymes was detected. From pure culture SRM capable of DNRA, it is known that sulfate is preferentially respired even in the presence of the thermodynamically more favorable electron acceptor nitrate and that expression of DNRA-related enzymes is only induced in the absence of sulfate, which acts as repressor (54).

Nevertheless, an involvement of Nbg-4 and related *dsrAB*-carrying *Nitrospirae* in anaerobic sulfur oxidation cannot be ruled out. For example, dense cell suspensions of the SRMs *Desulfovibrio desulfuricans* and *Desulfobulbus propionicus* are capable of coupling sulfide oxidation to nitrate reduction (55) and S^0^ oxidation to electron transfer to a graphite electrode (56), respectively. In addition, *Desulfurivibrio alkaliphilus* was recently shown to grow by sulfide oxidation coupled to DNRA while encoding and transcribing *dsrAB* affiliated to the phylogenetic branch of reductively operating sulfite reductases (40). *D. alkaliphilus* encoded and expressed also all other genes of the canonical pathways of sulfate reduction while oxidizing sulfide coupled to DNRA. At the same time, it lacked all typical sulfur metabolism genes of chemolithotrophic sulfur oxidizers with the exception of a membrane-bound sulfide-quinone oxidoreductase (Sqr). This led to the proposal that the canonical pathway of sulfate reduction could act in reverse when coupled to Sqr (40). Interestingly, Nbg-4 encoded Sqr as well, which showed a moderate similarity (54% amino acid identity) to Sqr of *D. alkaliphilus*. However, Sqr of Nbg-4 could not be identified to be expressed in the analyzed rice paddy metaproteomes (Table S2). The overall picture is further complicated by the phylogenetic placement of Nbg-4 and related *dsrAB*-carrying *Nitrospirae* between the genus *Thermodesulfovibrio*, which contains exclusively sulfate-reducing species, and magnetotactic *dsrAB*-carrying *Nitrospirae*, which are proposed to be capable of sulfur oxidation. Since genes encoding the biosynthesis of 12 magnetosomes were not detected in the largely recovered genome of Nbg-4 and it was significantly more abundant in the completely anoxic bulk soil (Figure 1), a lifestyle comparable to magnetotactic *Nitrospirae* can be most likely excluded.

In a preceding study, exclusively members of the Deltaproteobacteria (*Syntrophobacter*, *Desulfovibrio*, unclassified Desulfobulbaceae, and unclassified Desulfobacteraceae species) were identified to respond by population increase towards higher sulfate availability in rice paddy soil (7). The current study utilized soil from exactly the same experiment and identified Nbg-4 as an additional potential SRM. Nbg-4 did clearly not respond by changes in population size towards sulfate availability (Figure 1) but most likely by a switch in energy metabolism, i.e., from nitrate reduction under sulfate-depleted conditions to sulfate reduction under sulfate-enriched conditions (see above). This interpretation is supported by porewater sulfate concentrations reported in the previous study (7), where sulfate concentrations steadily declined from 2.6 to 0.5 mM throughout the incubation period in gypsum-amended bulk soil but were below the detection limit in unamended bulk soil. Together, both studies reveal that rice paddy SRM may follow different ecological strategies, either by activity response coupled to growth (Deltaproteobacteria) or by switching the energy metabolism to maintain a stable population (Nbg-4). Interestingly, species-level OTUs obtained in the previous study and which fall into a phylogenetic lineage resembling the Nbg-4 cluster (Fig. S2), constituted relative population sizes of up to 0.2% of the overall bacterial community in bulk soil irrespective of gypsum treatment (re-analyzed from 7). As such, these novel *Nitrospirae* constitute moderately abundant members of the bacterial bulk soil community. This is in accordance to a study of three different Chinese rice paddy soils, where comparable population sizes were recorded (52).

Nbg-4 and related *dsrAB*-carrying *Nitrospirae*, which were all recovered from groundwater systems, clearly formed a separate lineage within the *Nitrospirae*. This was supported by three independent phylogeny inference approaches as based on highly conserved marker genes, the *dsrAB* genes, and the 23S rRNA gene (Figure 4). Further indirect evidence was provided by the same branching pattern of 16S rRNA genes affiliated to the phylum *Nitrospirae* and recovered from the same microcosms (Fig. S2). In accordance with the performed gAAI analysis, Nbg-4 and related *dsrAB*-carrying *Nitrospirae* that form this separate lineage, constitute a newly discovered genus (Table S5). In addition, Nbg-4 represents a clearly distinct species in comparison to all members within this novel genus as based on the performed gANI analysis (Table S4). Based on its distinct potential physiology, separation into an own phylogenetic lineage, and predominant occurrence in habitats of moderate temperature, the following name is proposed for Nbg-4: *Candidatus* Sulfobium mesophilum [etymology: *Sulfobium* gen. nov. (Sul.fo'bi.um. L. n. *sulfur* sulfur; Gr. n. *bios* life; N.L. neut. n. *Sulfobium* a living entity metabolizing sulfur compounds), *S. mesophilum* sp. nov. (me.so'phi.lum. Gr. adj. *mesos* middle; Gr. adj. *philos* friend, loving; N.L. neut. n. *mesophilum*, loving medium temperatures)].

## Materials and methods

### Rice paddy microcosms

Soil from planted rice paddy microcosms described in S. Wörner et al. (7) was analyzed. In brief, microcosms were sampled destructively after 58-59 days of a greenhouse incubation to obtain rhizosphere and bulk soil of microcosms treated without (control) and with gypsum (0.15% (w/w) CaSO_4_×2H_2_O). In addition, freshly flooded soil was incubated for three days in the absence of a rice seedling and denoted as T_0_. As such, the experimental setup resulted in five different soil habitats: bulk soil with and without gypsum addition, rhizosphere soil with and without gypsum addition, and freshly flooded soil. Sampling from the different soil compartments and DNA extraction based on beat beating and phenol-chloroform extraction were as described in S. Wörner et al. (7).

### Metagenome sequencing, assembly, and binning

Rhizosphere- and bulk soil-derived DNA extracts were obtained from four separate microcosms per treatment (gypsum and control). In addition, three DNA samples were obtained from freshly flooded soil. For each replicate, 2 µg of DNA were used for metagenomic library preparation and paired-end sequencing (2 × 100 bp) on an Illumina HiSeq 2000 platform at the King Abdullah University of Science and Technology, Thuwal, Saudi Arabia. Raw reads were processed in the CLC Genomics Workbench 5.5.1 (CLC bio, Aarhus, Denmark) using only paired-end reads >50 bp with ≤1 ambiguity and a quality score ≥0.03 (corresponds to 99% accuracy). *De novo* assembly of pooled reads per habitat type was done in CLC using a k-mer size of 41 (determined as optimal in preliminary tests). Contigs with <2000 bp were discarded. Scaffolds containing 16S rRNA genes, 23S rRNA genes, or *dsrAB* were identified by a blastn search (57) against the respective SILVA reference databases v.123 (58) or a *dsrAB* reference database (37). Coverage of scaffolds was determined in CLC using 100% identity over the full length of quality trimmed reads. This was done for each sequenced replicate separately for statistical analysis and in addition using pooled replicates per habitat type for genome binning.

Genome binning was performed according to M. Albertsen et al. (59) using the gypsum and control treatment as differential coverage conditions (Fig. S3). From the 159 obtained genome bins, a *dsrAB-*carrying *Nitrospirae* bin assembled from gypsum-treated bulk soil was selected for further refinement (Figure S1). First, quality-trimmed reads that mapped to the *Nitrospirae* bin as well as to taxonomically unaffiliated scaffolds of similar coverage were re-assembled in CLC and binned as outlined above. Thereafter, obtained scaffolds were co-assembled with quality-trimmed reads of the first step using SPAdes (60). Binning resulted in the genome bin Nbg-4 (***N****itrospirae* genome bin from **b**ulk soil treated with **g**ypsum). Using this procedure, the genome of Nbg-4 could be extended from 1.15 Mbp with 57 out of 107 queried essential single-copy genes (ESG) to 2.77 Mbp that covered 92 ESGs, with 91 of these ESGs being present as one copy. Assembly refinement of a 23S rRNA gene fragment encoded at the end of one Nbg-4 scaffold is described in Supplementary Information. Completeness, contamination and strain heterogeneity of Nbg-4 were evaluated using CheckM (61). To assess its relative abundance in the different soil habitats, quality-trimmed reads of sequenced soil replicates were mapped with a similarity threshold of 100% over the complete read to 15 the Nbg-4 scaffolds using CLC. Mapped reads were normalized to RPKM values (reads per kilobase of scaffold per million reads).

### Annotation and additional analyses

The MicroScope platform was used for automatic annotation (62, 63). Annotation refinement was done as follows: proteins with an amino acid identity ≥40% (over ≥80% of the sequence) to a SwissProt entry (64) were annotated as homologous to proteins with a known function. Proteins with an amino acid identity ≥25% (over ≥80% of the sequence) to a SwissProt or TrEMBL (64) entry were annotated as putative homologs of the respective database entries.

Genome-wide average nucleotide identity (ANI, 47) and average amino acid identity (AAI, 45) comparisons were performed using the web service of the Konstantinidis laboratory at the Georgia Institute of Technology, GA, USA (enve-omics.ce.gatech.edu). The index of replication (iRep) was calculated using the iRep software (38). SAM files needed as input for iRep were created using bowtie2 (65).

To estimate the effect of soil habitat, gypsum treatment and the interaction thereof on the relative abundance of the *Nitrospirae* genome bin, a two-way ANOVA was performed based on RPKM values of its longest scaffold (106,945 bp) in the different replicated metagenomes. This was done using the base package of the program R, version 3.1.1 (66). Assumptions of variance homogeneity and normality were tested using Levene’s test in the R package lawstat (67). Significant differences between differently treated soil habitat types were inferred using Tukey’s test of honest significant difference.

### Metaproteomics of rice paddy soils

Total proteins were extracted from the same replicated soil samples as used for metagenome sequencing. Protein extraction and subsequent in-gel tryptic digestion followed the procedure outlined in R. Starke et al. (68). Briefly, 2 g of soil was used for a phenol extraction procedure with a subsequent ammonium acetate precipitation. Tryptic peptides were analyzed using a UPLC-LTQ Orbitrap Velos LC-MS/MS (69). Peptide searches were performed using the MaxQuant algorithm with the following parameters: tryptic cleavage with maximum two missed cleavages, a peptide tolerance threshold of ±10 ppm and an MS/MS tolerance threshold of ±0.5 Da, and carbamido methylation at cysteines as static and oxidation of methionines as variable modifications. As sample specific database, the Nbg-4 genome was used. Proteins were considered as identified with at least one unique peptide with high confidence (false discovery rate-corrected p-value <0.01). To check for false positive assignments, selected metaproteome replicates were also searched against the complete bacterial protein database of NCBI (08/2017).

### Phylogenetic analysis

Additional *Nitrospirae* genome bins carrying *dsrAB* were identified using a blast search (57) against NCBI’s sequence repositories (70). Only *Nitrospirae* genome bins with a completeness above 70% and a contamination below 5% according to CheckM (61) were considered for further analysis. The phylogenetic affiliation of Nbg-4 and public *dsrAB*-carrying *Nitrospirae* genome bins was inferred using a phylogenomics approach based on 43 conserved marker genes with largely congruent phylogenetic histories as defined by D. H. Parks et al. (61) as well as using *dsrAB* and 23S rRNA genes as phylogenetic markers. Respective maximum likelihood trees were calculated using RAxML v8.2.9 (71) as implemented on the CIPRES webserver (72, www.phylo.org). Details are provided in Supplementary Information.

### Sequence information

All sequences are available in the Short Read Archive of NCBI under bioproject number PRJNA391190. The draft genome of Nbg-4 has been deposited in EMBL under the study accession number PRJEB21584. The mass spectrometry data have been deposited to the ProteomeXchange Consortium via the PRIDE partner repository (73) with the dataset identifier PXD007817.

## Acknowledgements

We are grateful to Prof. Dr. Bernhard Schink and Dr. Nicolai Müller for helpful discussions and support in naming the novel *Candidatus* genus and species.

## Supplementary figure legends

**Figure S1.** Phylogeny of deduced DsrAB sequences of *Nitrospirae* bacterium Nbg-4 and related *dsrAB*-carrying *Nitrospirae* bacteria recovered from metagenomes of groundwater systems (42, 43). A maximum likelihood tree were inferred using the RAxML algorithm (71). Bootstrap support is indicated by closed (≥90%) and open (≥70%) circles at the respective branching points. *Nitrospirae* bacteria with *dsrAB* that underwent horizontal gene transfer are marked with an asterisk. The scale bar indicates 10% estimated sequence divergence.

**Figure S2.** Maximum likelihood 16S rRNA gene tree showing the phylogenetic position of species-level OTUs affiliated to the phylum *Nitrospirae*, which were obtained in a previous study (7) using the same rice paddy soil samples as analyzed in the current study. The tree was reconstructed using the RAxML algorithm (71) as implemented in ARB (74) using 1,222 unambiguously aligned nucleotide positions and a 50% conservation filter for the domain Bacteria. The representative 454 amplicon sequences were added to the tree by using ARB’s Parsimony Interactive tool as indicated by the dashed branch. Solid circles indicate ≥90% and open circles ≥70% bootstrap support (1000 replications). The bar represents 10% inferred sequence divergence.

**Figure S3.** Schematic overview of the bioinformatics workflow to obtain the high quality draft genome of *Nitrospirae* bacterium Nbg-4.

## Supplementary table legends

**Table S1.** Key characteristics of sequenced metagenomes.

**Table S2**. Annotation and locus of genes involved in energy and biosynthesis metabolism in *Nitrospirae* bacterium Ngb-4. Expression of respective genes as proteins is indicated in the metaproteomes of the respective analyzed soil replicates.

**Table S3.** Main characteristics of members of the phylum *Nitrospirae*. 34

**Table S4.** Genome-wide average nucleotide identity (gANI) of *Nitrospirae* bacterium Ngb-4 in comparison to other members of the phylum *Nitrospirae*.

**Table S5.** Genome-wide average amino acid identity (gAAI) of *Nitrospirae* bacterium Nbg-4 in comparison to other members of the phylum *Nitrospirae*.

